# Antiviral activity of the hemolymph of *Podalia sp and M. albicolis* (*Lepidoptera: Megalopigydae*)

**DOI:** 10.1101/2022.05.25.493399

**Authors:** N.D. Carvalho, S.P. Curti, M.I. Oliveira, H.K. Rofatto, C.A. Figueiredo, K Senna de Villar, R.F. Magnelli, R.Z. Mendonça

**Author notes:** **Corresponding author:** Ronaldo Zucatelli Mendonça. Laboratório de Parasitologia, Instituto Butantan, Av. Vital Brasil 1500, São Paulo, Brasil. CEP:05503-000.

## Abstract

Potent antiviral activity against measles, influenza, picornavirus and herpes simplex viruses was observed in the hemolymph of *Podalia sp* and *M. Albicolis* (*Lepidoptera: Megalopigydae*). The antiviral proteins responsible for this activity were isolated by gel filtration chromatography using a gel filtration column system (Superdex 75) and further fractionated using a Resource-Q ion exchange column system. Experiments with the semi-purified protein led to a 128-fold reduction in picornavirus production, 64-fold reduction in measles virus production and a 32-fold reduction in influenza virus replication. qPCR showed a significantly lower level of herpes virus transcription. In addition no citotoxicity and genotoxicity effect was observed for Vero cells, suggesting a very interesting potential antiviral activity.

## 1. Introduction

Viral infections cause a significant number of human diseases. While vaccine efforts have proven successful for preventing and eradicating some viral infections, many viruses can not be targeted by immunization. Other form of control include the use of antiviral drugs; Meantime, there are currently few licensed and efficacious drugs available for prophylactic and therapeutic antiviral treatments. Invertebrates A likely source of antivirals are the invertebrates. They do not have a developed immune system as in mammals, but the hemolymph of these artropods contains substances with potent antibacterial action (Amer et al, 2019., Oliveira et al, 2019., Díaz-Roa et al, 2019., Diniz et al, 2018, Díaz-Roa et al, 2018, Abreu, et al, 2017, Candido-Ferreira et al, 2017, Chaparro and Junior, 2016, Sayegh et al, 2016., Pavillard and Wright, 1957), virus (Olicard et al, 2005a., Olicard et al, 2005b., Dang et al, 2006, 2011, Mondotte et al, 2018., Cociancich et al, 1994., Ferrandon et al, 1998, Vistnes et al, 1981) fungal (Lamberty et al, 1999, Lauth et al, 1998., Fehlbaum, et al 1994 or parasitoid, infestations (Whitman et al 2019., Lacerda et al, 2016., Marr et al, 2012., Bell, 2011, Rangel et al, 2011, Gao et al, 2010, Fieck et al, 2010., Pulido et al, 2008. In invertebrates, the protector responses are based in the peptide with antimicrobial action (Imuler and Hoffmann, 2000; Krutzik et al., 2001; Underkill and Orinsky, 2002), hemolymph coagulation (Iwanaga et al., 1978), melanin formation (Sugumaran, 2002), and lectin-mediated complement activation (Fujita, 2002). In addition to these enzyme cascades, a variety of agglutinin-lectins and reactive oxygen producing and phagocytic systems cooperate with immune reactions to kill invading pathogens (Bogdan et al., 2000). Antiviral activity from insect hemolymph have also been described by Chernysh et al., 2002, Greco et al., 2009., Carvalho et al., 2017. However, antiviral defense and any interactions between antiviral and other antimicrobial defenses remain unexplored (Popham et al., 2004). Similarly, the phenol oxidase, an enzyme obtained from the tobacco budworm (*Heliothis virescens*) hemolymph, was reported to exhibit antiviral activity against several vertebrate viruses in vitro (herpes simplex viruses types 1 and 2, vesicular stomatitis virus, human parainfluenzavirus 3, coxsackievirus B3 and Sindbis virus) (Ourth and Renis, 1993). Earlier, we have demonstrated the presence of pharmacologically active substances in the hemolymph of *Lonomia obliqua* with positive effect on viability and replication cell culture (Mendonça et al., 2008; Raffoul et al., 2005; Souza et al., 2005; Maranga et al., 2003) and on recombinant protein production (Mendonça et al., 2008. Carmo et al, 2012) describes a system for the protein expression in Sf9/baculovirus cells using the recombinant DNA to obtain a protein from the L. obliqua caterpillar that displays a potent antiviral action obtained by Greco et al, 2009. Recently, our group has studied the presence of antiviral agents in diferents animals source (Lima-Netto et al, 2012., Carmo et al, 2015, Toledo-Piza et al, 2016, 2018; Carvalho et al, 2017; Coelho et al, 2015, 2018). So, the objectives of this work were the identification and isolation of antiviral substances in hemolymph from *Podalia sp and M. Albicolis* (*Lepidoptera: Megalopigydae*)

## 2. Methods

### 2.1. Cells

Vero cell lines (African green monkey kidney-ATCC CCL-81), MDCK and L-929 cells were grown in 25cm^2^ plastic cell culture flasks or in multiwell plates using Leibovitz-15 (L15) medium containing 0.9 g L-1 of D-galactose, 0.3 g L-1 of L-glutamine and supplemented with 5% fetal bovine serum (FBS). Viable cell counts were performed in Neubauer chamber using Trypan blue (0.05%) exclusion.

### 2.2. Hemolymph collection

The hemolymph of *Podalia sp and M. Albicolis* were collected from larvae after setae had been cut off. The collected hemolymph were clarified by centrifugation at 1000×g for 10min and filtered through a 0.2μm membrane and stored at 4 °C.

### 2.3. Hemolymph fractionation by chromatography

After centrifugation and filtration, 1mL of hemolymph of *Podalia sp or M. Albicolis* were loaded on a gel filtration chromatography system equipped with a Superdex 75 column (Amersham Pharmacia Biotech) and eluted with sodium phosphate buffer at 0.5mL/min. The elution was monitored at 280 nm and 0.5mL fractions were collected. The fractions were then analysed by SDS-PAGE electrophoresis.

### 2.4. Determination of the cytotoxic effects

The cell viability was determined by MTT assay. To this, Vero cells were cultured in 96-well plates in Leibovitz medium supplemented with 5% of fetal bovine serum. After 48 h, HMG with concentrations ranging from 10mg/mL to 0.1mg/mL was added to the wells culture medium. After 24 hours of exposure the supernatant was discarded and MTT at concentration of 500 microgram/mL was added to the growth medium for 4 hours. After this, the culture medium was removed and 100mL of DMSO was added. After this, the plates were shaken for 30 minutes and read on a spectrophotometer at 570 nm.

### 2.5. Comet test on Vero cells

For analysis of possible genotoxicity caused by the hemolymph of *Podalia sp, or M. Albicolis* cells, Vero were treated with 1% v / v of total hemolymph or its fractions, for 24 hours.

The general procedures used in the alkaline comet test were performed as described by Singh et al., 1988. The experiments were performed in the dark to minimize any DNA damage caused by ambient light. Each preparation was photographed at a resolution of 1376 x 1032 pixels and 100 cells from each group were analyzed. Quantitative DNA damage was determined by the increase length and intensity of the comet tail corresponding to each cell.

The classification was made according to the size of the tail into 4 classes> 0 = undamaged; 1 = little damage; 2 = moderate damage and 3 = full damage (Savi, 2004). Comets with small or nonexistent heads and broad and diffuse tail were not included in the analysis as described by Bauer et al., 1998 and Schnurstein and Braunbeck, 2001).

### 2.6. Virus and cell infection

Measles virus (Edmonston strain), Influenza virus (H1N1), herpes Simplex Virus (HSV-1) and picornavirus (EMC-encephalomyocarditis virus) were used to determine the antiviral activity of the hemolymph. To this, MDCK cells were infected with 0.1 moi (multiplicity of infection) of influenza virus, Vero cells were infected with 0.01 m.o.i of measle or herpes virus and L-929 cells were infected with 0.001 m.o.i of EMC virus.

### 2.7. Determination of antiviral effect

To determine the antiviral effect, 96 well plates, with the different cells, were treated or not with 1% v/v of total hemolymph (*Podalia sp or M Albicolis*) or with 0.05 mg/mL of its fractions, 1 h prior infection. After this, the medium were removed and cell culture were infected with 100μL of serial virus dilution (r:2) of different virus, in quadruplicates. After 1 hour adsorption at room temperature, the medium was removed and the plate replace with medium with 1% FBS. Infected and uninfected cultures, without hemolymph treatment, were prepared as negative and positive controls. All culture were incubated at 33°C. Plate cultures were observed daily for cytopathic effect (CPE) determination as described by Griffiths and Thornton (1982). Virus titers were determined by monitoring the CPE and the endpoint dilution were determined as the highest dilution of virus able to induce CPE in 50% of cells). The reduction of virus infection was determined by diference between control infected culture and the culture infected and treated with total hemolymph or its fractions. After cytopathic effect determination, the culture medium was removed and the cells in the plate were stained with crystal violet (0.2% in 20% ethanol).

### 2.8. Determination of antiviral activity of *Podalia sp* and *M. albicolis* against infected cells with influenza virus, using fluorescent antibodies

The protector effect of hemolymph was also determined by immunofluorescence assay. To this, MDCK cells were cultured in 96-well microplates. One hour before infection by 0,1 MOI of Influenza virus (H1N1), cells were pre-treated with 1% (v/v) total hemolymphm of *Podalia sp or M. albicolis* and maintained at 33°C. After 72 hours, the cells were rinsed twice with PBS and fixed in 4% paraformaldehyde (Fluka) for 20 min. After fixation, the cells were washed three times with PBS, permeabilized with Triton X-100 0.1% (Sigma) for 10 min and blocked with PBS, 1% BSA (Sigma) and 50 μg/mL RNAase A (Invitrogen) at 37 °C for 1 h. After this, the cells were incubated with 10 μg/ mL of mouse monoclonal anti-Influenza FITC antibody (Millipore) and with in blue evans stain, in 50 μL of PBS+1%BSA solution at 4 °C overnight. The cells were washed three times with PBS containing 0.05% Tween 20 (Sigma) before incubations with 20 μg/mL propidium iodide (Sigma) in 50 μL of PBS + 1% BSA for 1 h. After this incubation, the cells were washed five times with PBS + Tween 20 and 50 μL of an anti-fading solution with PBS, 50% glycerol (GE Healthcare) and ρ-phenylenediamine 0.1% (Sigma) was added to each well. The images were acquired with an confocal microscope (LSM 510 META, ZEISS)

### 2.8. Quantitative real-time PCR assay of cells infected by herpes virus

Total RNA was extracted from 200μL of VERO cells culture infected by herpes virus (treated or not with hemolymph) using the MagNA Pure extractor (Roche, Basel, Switzerland). The Herpes simplex virus quantitative real time PCR assay was performed was described by Read et al., ^16^. For the real time PCR, the 20μl reaction contained 12μl of the SYBR® Green PCR Master Mix (Applied Biosystems, Foster City, CA EUA), 10μM of each primer, 6.5μl H2O, and 5 μl cDNA. Cycling conditions were as follows: 94°C for 5 min, followed by 35 cycles of 94°C for 30 s, 57°C for 20s and 72°C for 40s. Four positive controls containing known copy numbers of transcribed virus RNA were run on each plate as quantification standards. A reaction mixture containing water as the template was run on each plate as negative control. The data were analyzed with SDS software (version 2.1 AB).

## 3. Results

### 3.1. Determination of the cytotoxicity of the hemolymph

In this study, the citotoxicity properties of the hemolymph of *Podalia sp* and *M. Albicolis* were examined in VERO cells by MTT assay. As can see in the **Figure 1**, hemolymph concentrations lower than 2% (v/v) were not toxic to VERO cells.

**Figure 1:**
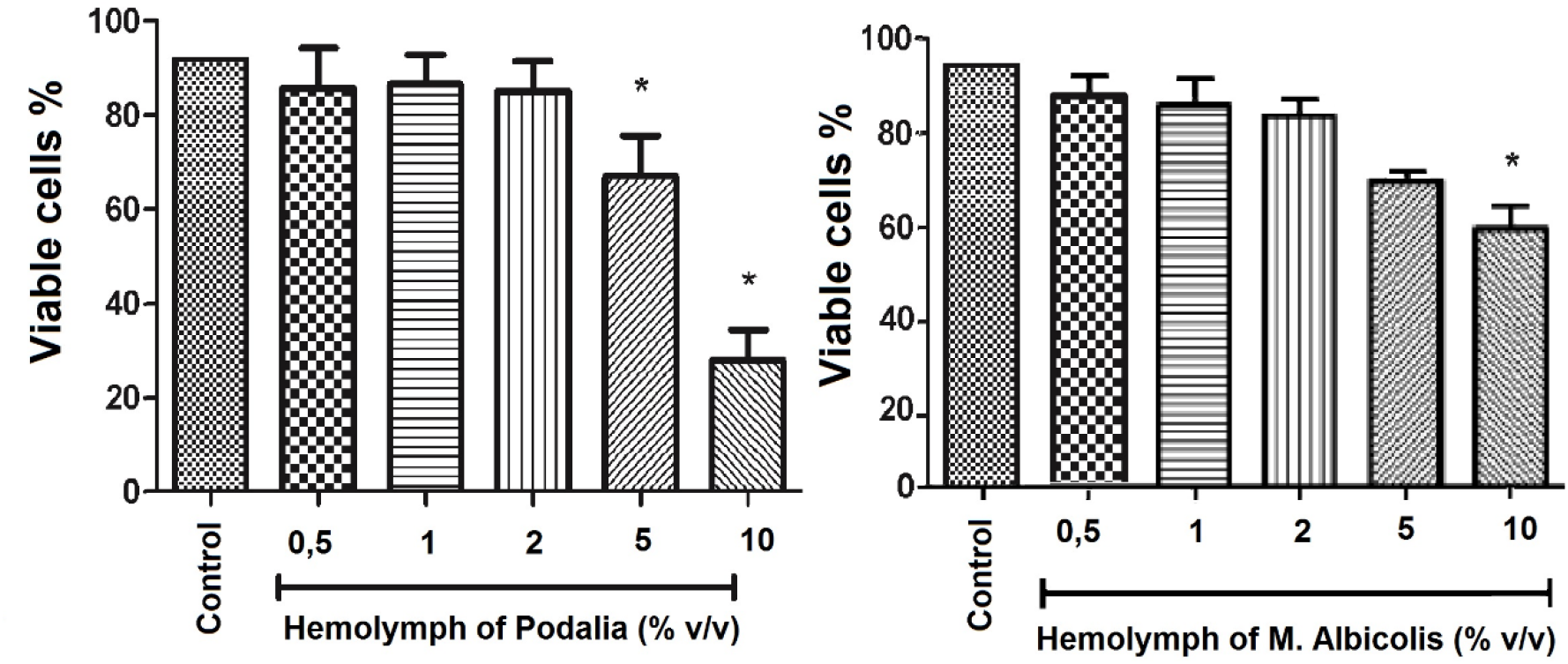
Effect of the hemolymph of *Podalia sp* and *M. albicolis* on the viability of VERO cells exposed to different amounts of hemolymph (0,5 to 10%v/v). After 96h of incubation with hemolymph the cell viability was evaluated by MTT reduction assay. Media of three experiments *p<0.05 compared to control (ANOVA and Dunnett’s Multiple Comparison Test).

### 3.2. Hemolymph fractionation by chromatography

After centrifugation and filtration, hemolymph of *Podalia sp* and *M. Albicolis* were loaded on a gel filtration chromatography system equipped with a Superdex 75 column. The results obtained are showed in **figure 2**. Different chromatography perfil were obtained. Fractions of each hemolymph were harvested and storage for further tests. All fractions collected were added to cell cultures for antiviral activity determination.

**Figura 2:**
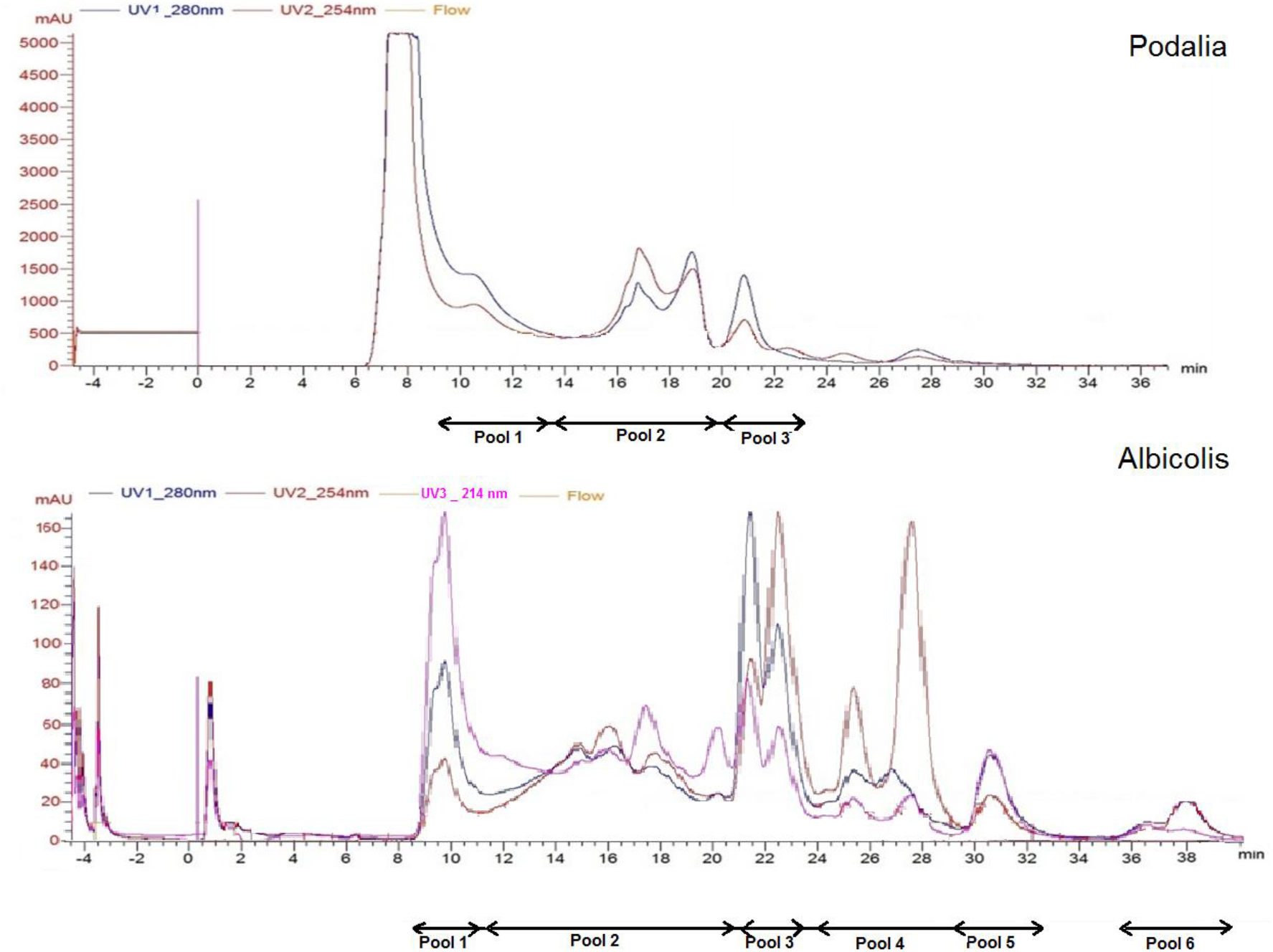
Purification of hemolymph from *Podalia sp* (A) or *M. albicolis* (b), by gel filtration chromatography on a “Superdex 75” column. Elution was performed with Tris (20mM) and Nacl (150mM) solution. The flow used was 0.5 ml / minute and fractions of 1 ml were collected. The fractions were separated into fractions as indicated by the bars.

### 3.3. Genotoxicity of the hemolymph of Podalia SP

The determination of the genotoxic potential of hemolymph was performed by the comet method. For this, the genotoxicity of various concentrations of fraction 3 was tested in VERO cells as described in ‘‘Materials and methods”. As positive control, the cells were treated with 200 μm of H202. As can be seen in **Figure 3**, no deleterical effect resulting from the treatment of the cells was observed. In our study, according to the classification, about 90% of comets were Class 0 (undamaged) or Class 1 (little damage) when a amount equal lower than 3% v/v was used

**Figure 3.**
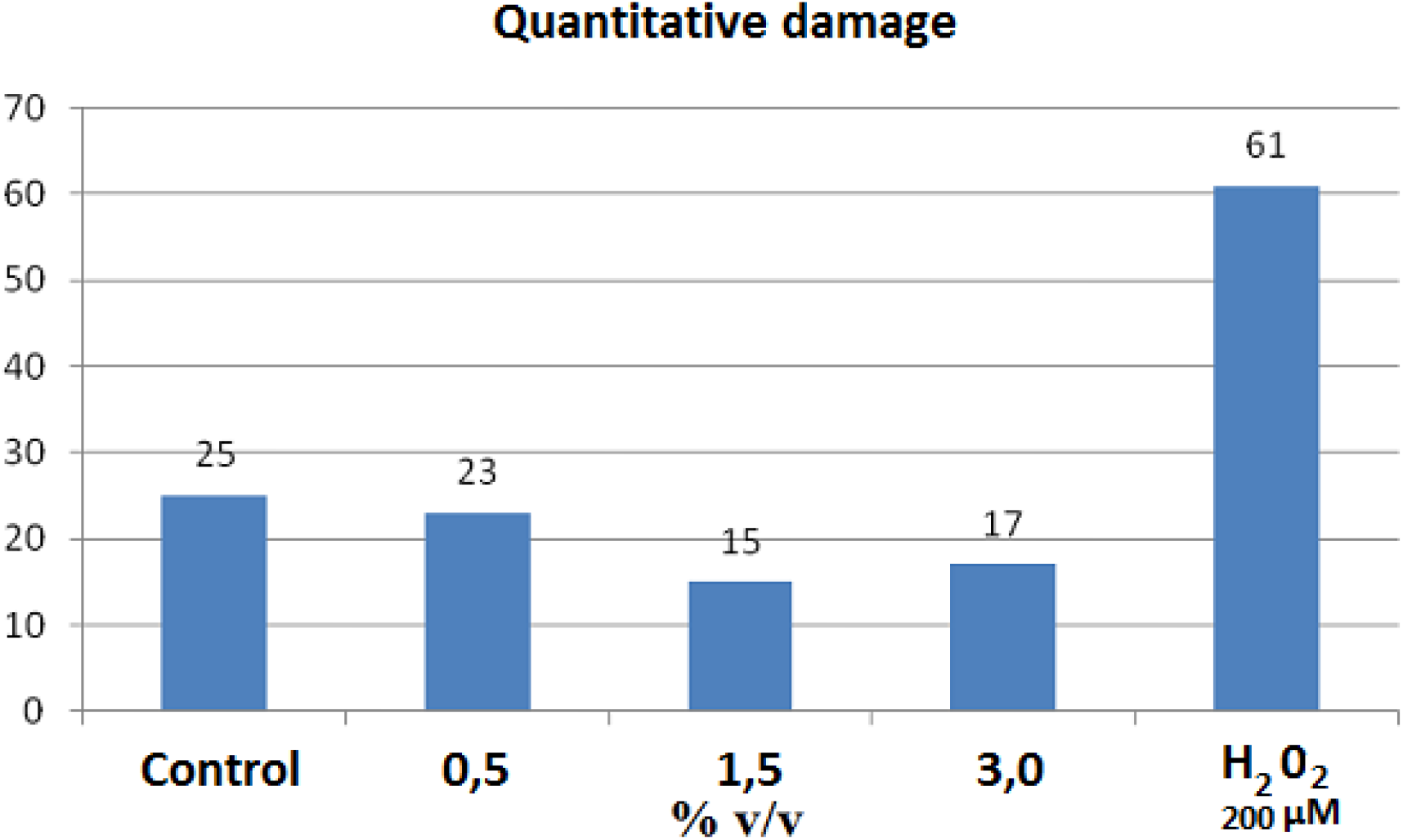
Genotoxicity of fraction 3 of *Podalia SP* in VERO cells. The cells were treated with 0.5; 1.5 and 3% (V/V) of the fraction 3, or with 200 μm of H202. A total of 100 comets/blades were randomly selected and analyzed using visual classification based on the amount of DNA in migration. The classification was performed according to the size of the tail in 4 classes and determined the quantitative damage.

### 3.4. Antiviral activity of the hemolymph on the replication of the picornavirus (EMC)

To verify the effect of hemolymph on picornavirus replication, cultures of L929, treated or not with total or with semi purified fractions of *Podalia* hemolymph, were infected with serial dilution (r:2) of picornavirus (initial titer of 1 x 10^4^ TCID50). In the culture infected, the hemolymph as able to reduce in 7 dilution the virus titer **(Figure 4a).** In the **Figure 4b**, in showed the cell aspect in dilution 1/1024, in infected and non infected. Nevertheless, when the same procedure was performed in culture treated with hemolymph, cytopathic effect (CPE) was observed only until dilution 1:8 to fraction 3, showing that hemolymph of *Podalia* sp was able to inhibit the virus replication by 128-fold. Nevertheless, when no infected culture was pre-treated with hemolymph, no adverse effect in the cell culture was observed, and it remained highly viable (data not showed).

**Figure 4:**
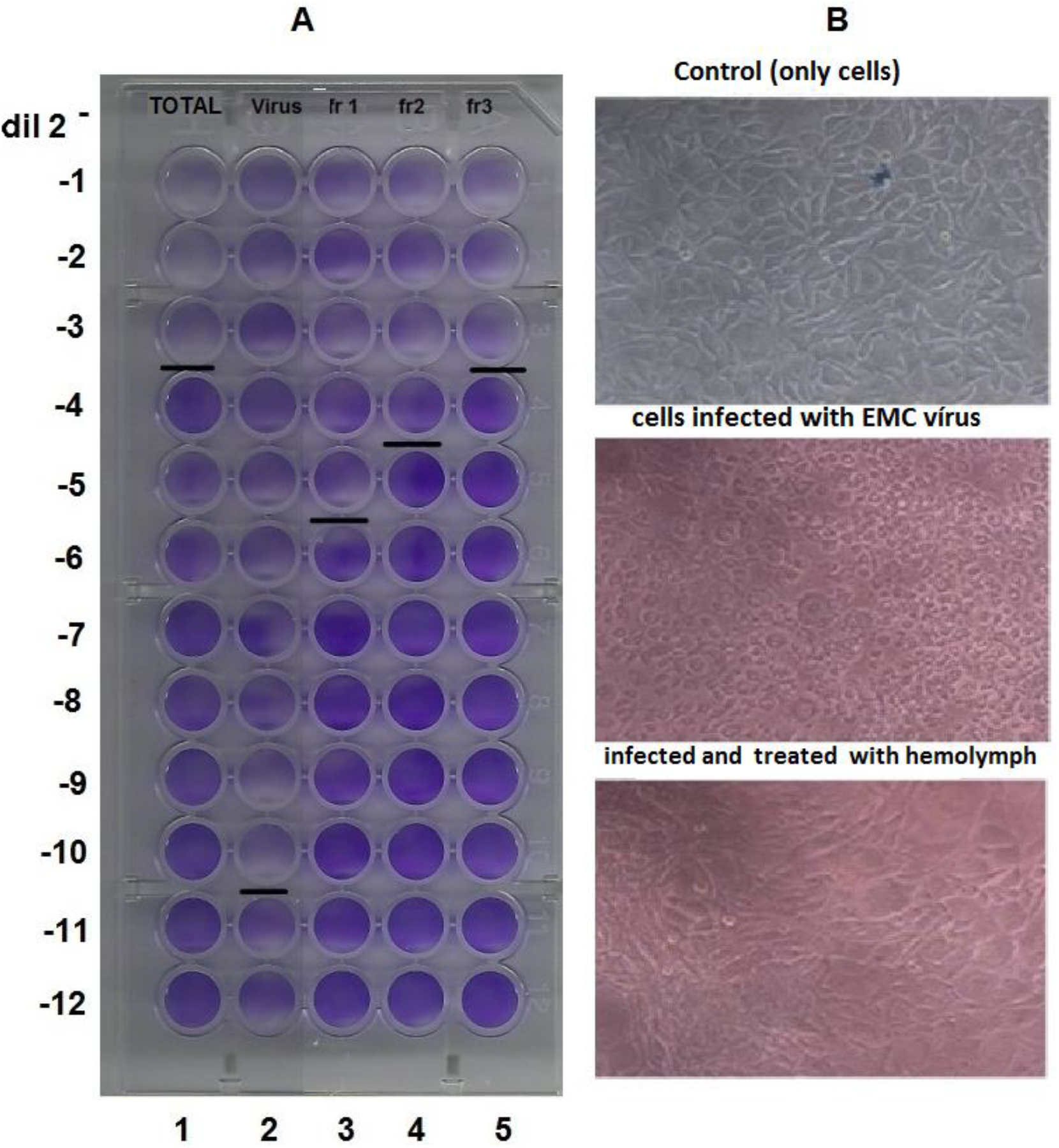
Effect of hemolymph of *Podalia sp* on virus replication. Cultures of L929, treated or not with 1% v/v of crude hemolymph or with 0.05 mg/mL of its semi purified fractions were infected with serial picornavirus dilution (r:2) (initial titer of 4 x 10^4^ TCID50). Cell cultures were observed after 72 h and the cytopathic effect determined on an inverted microscope. (A) (1) Infected and treated with hemolymph total; (2) *vírus control* (3) fraction 1; (4) fraction 2; (5) fraction 3. Cells without cytophatic effect are stained in blue. The last virus dilution seen as positive is marked with a black line.

### 3.5. Antiviral activity of the hemolymph on the replication of the measles

The same routine above was followed as in the experiment with the Measles vírus. After 96 h post-infection with measles, a sample was taken from each culture and titrated on 96-well plates to determine virus titers. When 1% hemolymph was used in the infected culture, a reduction of virus replication was 128 folder higher than observed in m control of infected culture **(Figure 5).**

**Figure 5:**
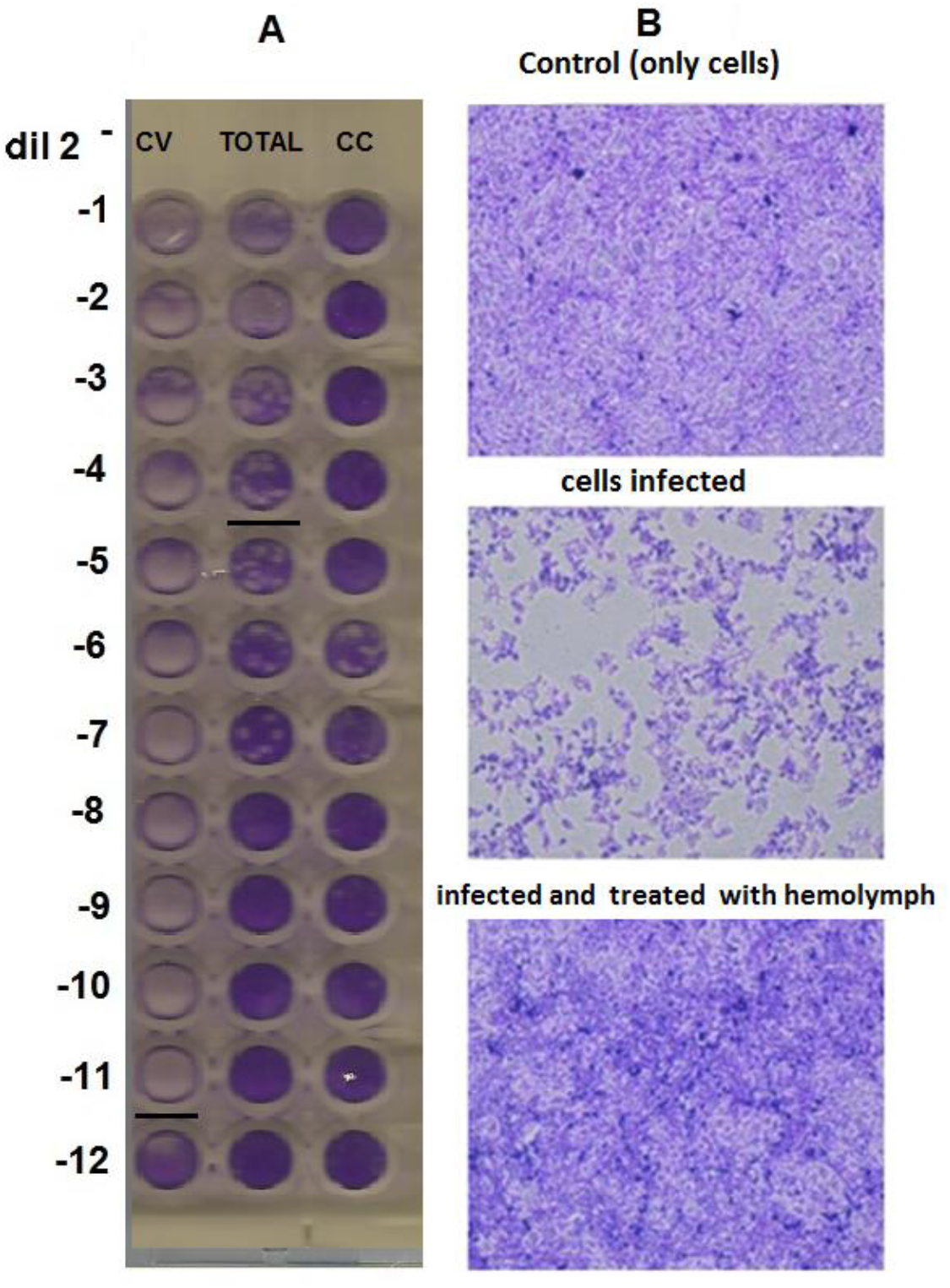
Antiviral effect of *Podalia sp* hemolymph. VERO cell culture treated or not with 1% v/v of total hemolymph from *Podalia sp* by 1 hour and then infected with 10^4^ DTCT/50 of Measles virus (Edmonston). After 96 hour, cultures were stained with crystal violet (0.2%) and observed under an inverted microscope (x40). (CC) Control cells; (CV) Infected cells (tHb) Infected cells and previously treated with total hemolymph

### 3.6. Antiviral activity of the hemolymph on the replication of the influenza

The same routine used above was followed to determine the effect of Podalia hemolymph against influenza virus (H_1_N_1_). MDCK cells pretreated with whole hemolymph of *Podalia sp* (1% v/v) or with 0.05 mg/mL of it fraction 3, were infected 1 hour after treatment with H_1_N_1_ vírus. As shown in **Figure 6**, the amount of virus produced in cultures supplemented with 1% whole total hemolymph and fraction 3 was approximately 32-fold lower than that obtained in the control infected culture. Cell cultures were observed after 72 h and the cytopathic effect determined on an inverted microscope **(Figure 6).**

**Figure 6:**
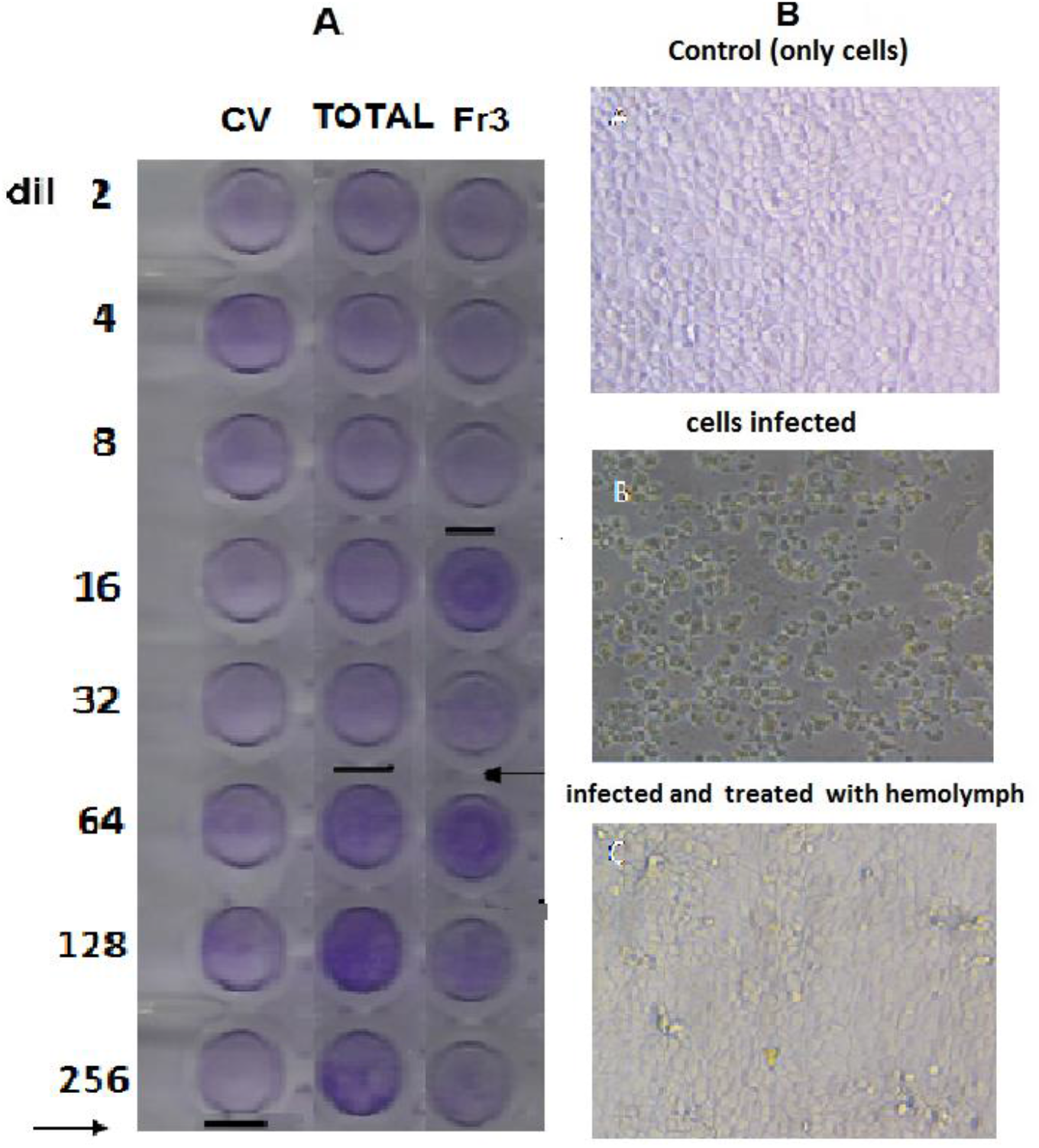
Antiviral effect of *Podalia sp* hemolymph against influenza virus. MDCK cell culture, treated or not with 1% v/v of total ou fraction 3 of *Podalia sp* hemolymph, was infected 1 hour after treatment with 10^3^ DTCT/50 of Influenza virus (H1N1). After 72 hour, cultures were stained with crystal violet (0.2%) and observed under an inverted microscope (x40). (Cv) Control of infected cells; (tH) Infected cells and previously treated with total hemolymph. (Fr 3) Infected cells and treated with fraction 3

### 3.7. Determination of antiviral action of *Podalia* e *M. albicolis* hemolymph by Imunofluoresce assay

To determine the antiviral effect of hemolymph against vírus da Influenza H_1_N_1_, MDCK cells were pretreated with hemolymph of *Podalia sp* or *M. Albicolis*. 1 hour after, cells were infected or not with Influenza vírus H_1_N_1_. Infected cells were observed after 72 hours of infection and then stained with specific anti Influenza A antibody (green) or Evans blue (red). Imunofluoresce of the cultures were observed by confocal microscopy. Can be observed at Figure 7, a significant reduction of virus focus can be observed to both hemolymph.

**Figure 7:**
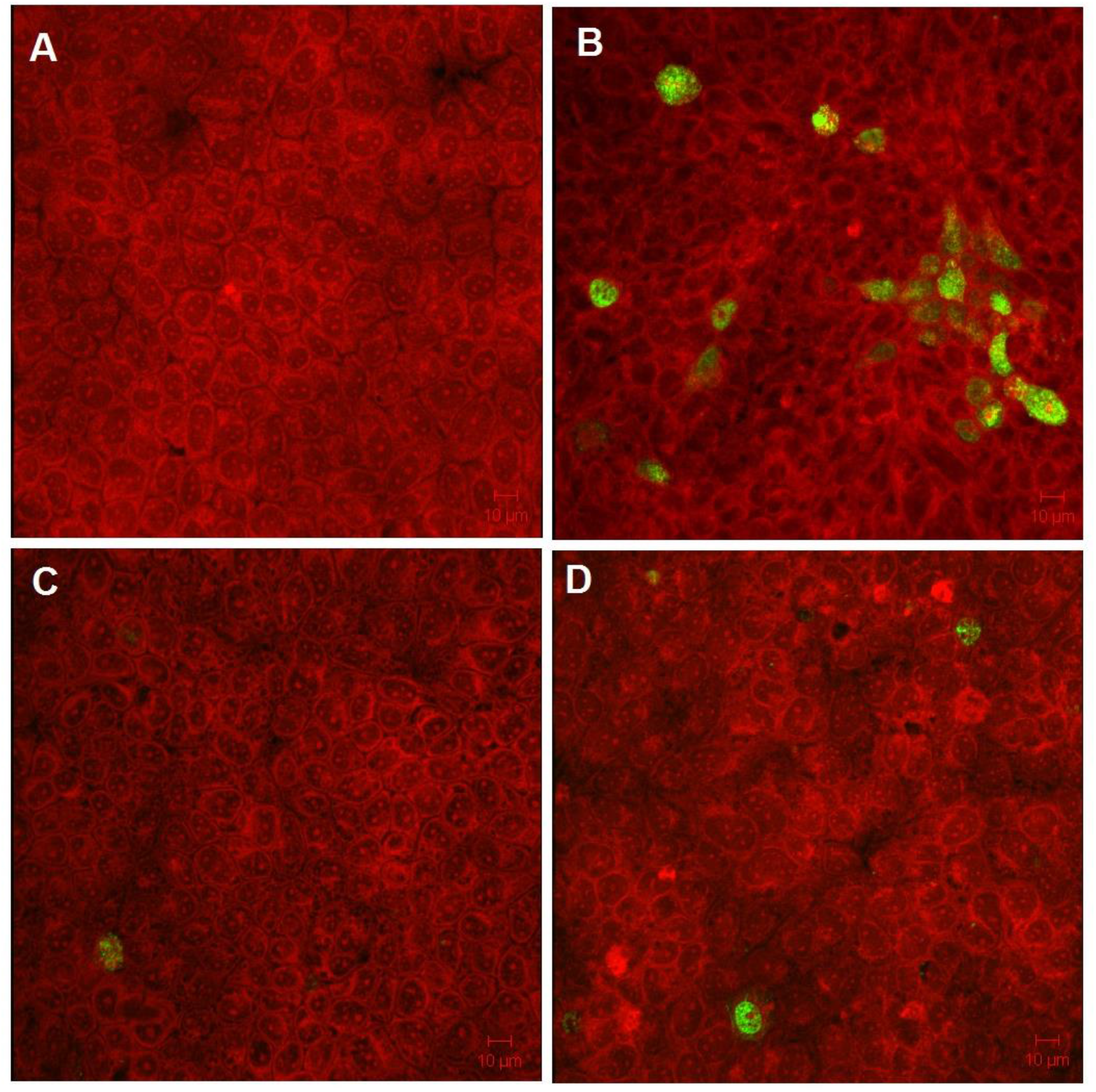
MDCK cells were cells were infecteds with Influenza vírus (H1N1).Cells were observed for 72 hour and then stained with antiinfluenza antibodies. A) Negative control (uninfected cells), B) cells infected with influenza virus C), cells infected and pre-treated with *Podalia sp* hemolymph D) cells infected and pre-treated with *M. albicolis* hemolymph. The cells were observed in confocal microscope and photographed. Green (anti influenza); red (Evans blue)

### 3.7. Determination of antiviral activity of the hemolymph *Podalia sp* e *M. albicolis* by qPCR

To identify any inhibitory effect of ***Podalia sp*** and ***M. Albicolis*** on vírus replication, the synthesis of herpes vírus was compared in infected cells, treated and not with fractions of *Podalia sp* and *M. Albicolis*. Total DNA extraction was performed at 96 hours after herpes infection, and the levels of intracellular herpes were measured. A standard curve for the amplification of both viruses was generated by a serial 10-fold dilution of the template to evaluate the efficiencies of both reactions. The resulting standard curves showed that the amplification efficiencies for the genes were between 95% and 100% with correlation coefficients greater than 0.99. The results of the qPCR showed a significantly increased level of herpes transcription in the inoculated cells at 4 days after viral infection compared to the mock. Was observed also that the total hemolymph and fraction 3 of the Podalia sp inhibit the replication of herpes virus **(Figure 8).**

**Figure 8:**
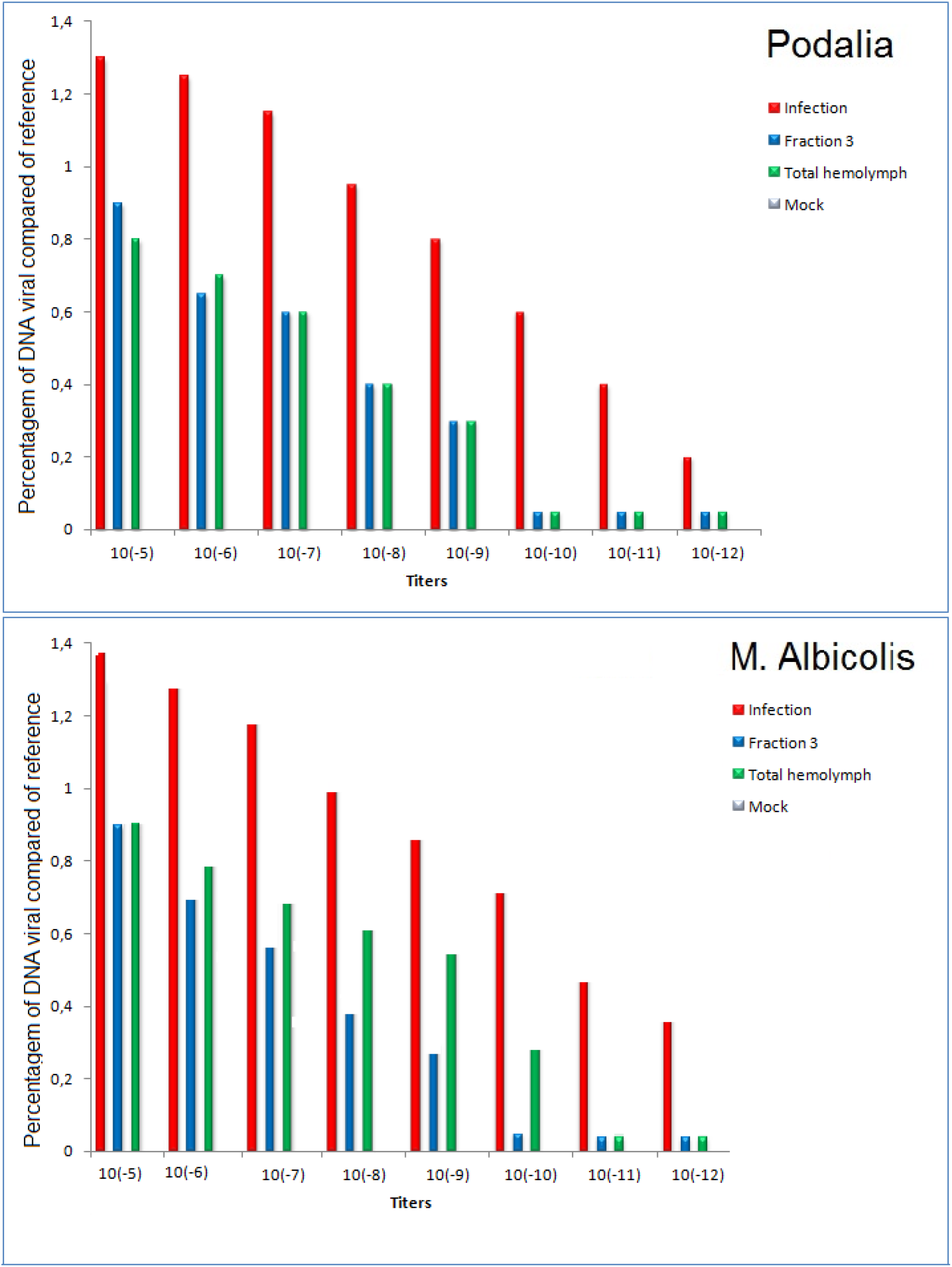
qPCR of herpes simplex viral DNA. Vero cells were infected by herpes virus (10^-5^ a 10^-12^) at the same time as treatment with fraction 3 and total of the *Podalia sp*. or *M. albicolis*. Total DNA was performed at 96 hs after infection. After reverse transcription, the levels of intracellular herpes simplex viral were measured by qPCR.

No reduction on vírus replication was observed witg others fractions. (data not showed).

## 4. Discussion

The cytotoxicity is caused by changes in cellular homeostasis that interfere with the capacity of the cells and their survival, their proliferation and the metabolic performance of their duties. Nardone 1977). In this study, it was found that the whole hemolymph of Podalia sp at concentrations below 5% and its purified fractions within the range of concentrations tested have no deleterious effect on VERO cells. Petricevich and Mendonça 2003, Karagöz, et al. 2003, Ourth, 2004 and Olicard et al. 2005 have carried out studies to identify substances with antiviral activity. These authors stress the importance of the identification of agents that show no toxicity to cells so that their application is feasible.

Various studies have reported the antiviral activity in products obtained from invertebrates. Cherrnysh et. al. (2002), have isolated two antiviral and antitumoral peptides from hemolymph of Calliphora vicina, which control viral infection when added before infection. Alloferon, one of the protein isolated was effective against Influenza vírus A e B. it due na intracelular action similar to interferon from vertebrate, inducing a intracellular response before the viralinfection (Chernysh et al., 2002). Olicard et. al (2005) showed that hemolymph of Crassostrea gigas inhibits HSV-1 replication in infected vero cell culture. The hemolymph of H. virescens larvae reduced the titers of the baculovirus HzSNPV (Popham et. al., 2004; Shelby et al 2006), as well of herpes simplex virus types 1 and 2, vesicular stomatitis virus, parainfluenza-3, coxsackie B3, Sindbis virus,and HIV-1 (Ourth and Renis 1993; Ourth, 2004). By the other hand, extracts of crustacean tissues have showed a broad spectrum of antiviral activity against enveloped and non-enveloped DNA and RNA viroses. This can be due to multiple inhibitors presente in the extracts (Pan et al., 2000). Peptides derived from scorpion venom showed virucidal activity against measles, SARS-CoV and influenza H5N1 viruses (Qiaoli Li et al., 2011) and HIV-1 (Chen et al., 2012). Dang et al., 2011, demonstrated that the total hemolymph and lipophilic extract of the digestive gland from abalone H. laevigata has activity against herpes simplex virus. Salas-Rojas et al., 2014, showed that the coelomic fluid of the sea urchin Tripneustes depressus has antiviral activity against Suid herpesvirus type 1 (SHV-1) and rabies virus (RV). A peptide derived from mastoporan has showed a broad-spectrum antiviral activity against enveloped viruses (Sample et al., 2013). Other peptide, isolated from a marine sponge Petromica citrine, has showed an antiviral activity of against bovine viral diarrhea vírus (Bastos et al., 2013). Lima-Netto et al., 2012, observed the antiviral effect of eggs wax from tick Amblyomma cajennense against influenza and picornavirus, for influenza virus, in this case, an amount as small as 12 μg/mL from a crude suspension of egg wax was able to reduce in 128 UHA (hemaglutinant unit) of influenza virus (H1N1). With picornavirus, this reduction was of 256-fold. Our group have purified an antiviral protein of approximately 20KDa from hemolymph of Lonomia obliqua; when added to cultures 1h before infection, this protein led to a 157-fold reduction in measles virus production and a 61-fold reduction in polio virus production (Greco et al., 2009). It has been suggested that it may act in the stages of the replication cycle of intracellular viruses, acting with a mechanism similar to that described for Alloferon. Recently, this antiviral protein was cloned and expressed and was able to inhibit the replication of picornavirus, rubella vírus (104 fold) and herpes simplex virus (106 fold) (Carmo, et al., 2012).

In this studie was tested hemolymph of Podalia sp for potential antiviral activity against picornavirus, influenza, measles, herpes simplex virus. Experiments with the semi-purified protein led to a 64-fold reduction in measles virus production, a 32-fold reduction in influenza and a 128-fold reduction in picornavirus production. The results of the qPCR showed a significantly reduction on level of herpes transcription.

The broad-spectrum antiviral activity of hemolymph of *Podalia sp* on various viruses, enveloped and non-enveloped DNA and RNA viruses, suggest an intracellular mechanism of action and that the protein could be a constitutive agent that acts on the innate immune system. This mechanism of action need further investigation. In this paper, we demonstrate the antiviral activity of Podalia sp hemolymph and its fraction 3 against herpes simplex (HSV-1). We show that the Podalia sp hemolymph and semi purified fraction 3 at concentrations below 88 μg / ml and 180 ng / ml, respectively. The tested concentration range have no deleterious effect on Vero and SIRC cells. In addition, the antiviral effect with pre-treatment of hemolymph cells was more intense than with hemolymph addition at the same time of infection or 1 hour after infection (data not shown). Cells treated prior to infection showed not only a reduction in viral titers, but also a decrease in the intensity of the cytopathic effect. HSV-1 and HSV-2 utilize various glycoprotein receptor complexes when entering host cells. Both express the gB, GC and GD, Geh and Gl glycoproteins. All except gC are essential for host cell entry, although the absence of gC reduces efficiency. gC makes the first contact with the host cell, the binding of heparin sulfate (HS) proteoglycans on the cell surface (Ashley and Corey, 1995; Bernard et al., 2007). In this study, it was found that hemolymph of both species exhibit HSV-1 inhibitory effect. Four probable mechanisms can be deduced regarding their antiviral of inflammatory cytokines induced by virus infection (Bernard et al., 2007). The outcome of HSV infections is dependent on a balance between virus spread and an effective immune response. Proper expression of IFNs, cytokines and chemokines is essential for efficient host defense against infection. Melchjorsen et al., 2009 showed that the initial response of HSV infected cells is secretion antiviral substances such as defensins and nitric oxide and cytokine production, including IFN and chemokines. Earlier we also have report that type I IFNs stimulate Nitric Oxide Production and Resistance to *Trypanosoma cruzi* Infection (Costa et al., 2006). As ocurr with trypanosomes, we believe that the mechanism of action of some antivirals is to stimulate cells to produce interferon, nitric oxide or other cytokines.

## Acknowledgements

This study was kindly supported by FAPESP 2016/24867-1 (Fundação de Apoio à Pesquisa do Estado de São Paulo).

